# Prevalence of sleep-disordered breathing and associations with malocclusion in children

**DOI:** 10.1101/560722

**Authors:** Maria Carlla Aroucha Lyra, Débora Aguiar, Mabel Paiva, Manuela Arnaud, Arnoldo Alencar Filho, Aronita Rosenblatt, Nicola Patricia Thérèse Innes, Mônica Vilela Heimer

**Affiliations:** Department of Pediatric Dentistry, University of Pernambuco, Camaragibe, Pernambuco, Brazil; School of Dentistry, University of Dundee, Park Place Dundee, DD14HR, Scotland

**Author notes:** Corresponding author (MC). Department of Pediatric Dentistry, University of Pernambuco, Camaragibe, Pernambuco, Brazil (Mônica Vilela Heimer). School of Dentistry, University of Dundee, Park Place Dundee, DD14HR, Scotland (Nicola Patricia Thérèse Innes). The author and co-authors contributed equally to this work (Maria Carlla Aroucha Lyra, Aronita Rosenblatt, Nicola Patricia Thérèse Innes, Mônica Vilela Heimer).

**Keywords:** Sleep-disordered Breathing, Child, Prevalence, Malocclusion, Body Weight

## Abstract

**Objective:** This study aimed to determine the prevalence of sleep-disordered breathing (SDB) and its association with malocclusion among children in Recife, Brazil.

**Methods:** 390 children aged seven to eight years took part in the study, comprised by the body mass measurement, orthodontic examination and parent’s information required by the Sleep Disturbance Scale for Children. Statistics tools were Pearson’s chi-square and Lemeshow test.

**Results:** SDB was found in 33.3% of the children and associated with overjet (p= 0.007), anterior open bite (p=0.008) and posterior crossbite (p= 0.001). There was no association between BMI and SDB. The multivariate logistic regression model indicated that the anterior open bite (p= 0.002) and posterior crossbite (p = 0.014) have an association with SDB.

**Conclusions:** Results of this study indicated that the prevalence of SDB was high and highly associated with malocclusion; anterior open bite and posterior crossbite are risk factors for SDB.

## INTRODUCTION

Scientific investigation on Sleep disorders usually reports to insomnia or excessive daytime sleepiness in adulthood and its negative impacts on normal physiological functioning and quality of life. However, very little is written on the prevalence of the condition in childhood, even though it impacts for 20 to 30% of problems reported by children’s health providers¹,². Inadequate sleep can lead to diverse comorbidities, such as delayed growth as well as cardiovascular, immunological and metabolic conditions, which can affect a child’s health and quality of life over the years³. Among the primary risk factors, the body mass index (BMI) is an important predictor of the clinical intensity of sleep disorders^3^.

Sleep-disordered breathing (SDB) includes a spectrum of upper airway disorders ranging from primary snoring (PS) to obstructive sleep apnea (OSA)^4^. PS is a respiratory noise, in which sleep architecture, alveolar ventilation, and blood oxygen saturation are maintained at normal values whereas OSA is partial or complete obstruction of the upper airways that impairs normal ventilation during sleep and normal sleep patterns^5^. The prevalence of SDB in children ranges from 0.7% to 16.9%; however, in the current literature, PS ranges from 4.3 to 16.9% and OSA from 0.7 to 3% ^6,7,8,9^.

Thus, abnormal craniofacial morphology, such as retrognathia and dental malocclusion contribute to upper airway obstruction and increase the risk of SDB^6,10^. Therefore, SDB is commonly associated with oro and dentofacial features and may be related to malocclusion. Failure to diagnose SDB in adults may have undesirable health outcomes^11^ which may require further education for both parents and providers of how occlusal abnormalities associates with SDB^12,13^.

Sleep-related issues often are not discussed during well-child visits, because of parents not raising the issue and primary care providers not asking about symptoms of sleep disorders. Sleep disorders are underdiagnosed in primary care practices^7^

Nevertheless, oral screening in children and the use of a validated questionnaire could improve early diagnosis and reduce the under-diagnosis of sleep disorders^14^.

This study aims to evaluate the prevalence of SDB and its relationship with malocclusion in Brazilian children.

## METHODS

A cross-sectional study, conducted with aged seven and eight year’s old children, attending local public schools in the city of Recife, Brazil.

The study was carried out following the Internal research Board of the University of Pernambuco by the record number CAAE 53308915.8.0000.5207 and followed the Resolution 466/2012 of National Commission of Ethics in Research and with the 1964 Helsinki declaration and its later amendments or comparable ethical standards, including Informed consent from all study participants. It also followed the STROBE protocol for observational studies (Additional file 1).

The sample size was calculated using the formula for cross-sectional studies in Epi Info Version 2000 (Atlanta, Georgia, USA), based on Schlarb et al.^1^, which is 20% prevalence rate for SDB, 95% confidence interval, 5% margin of error, a 1.5 design effect and the addition of 20% to make up for possible dropouts, a total of 435 children. A two-stage sampling method allowed the representativeness of the sample. The first stage was the randomization of public schools in each administrative district. In the second stage, classes were assigned by random computer number and all children attending those classes were invited to take part in the study.

The physical assessment comprised the measurement of body mass index (BMI), based on height (measured using a stadiometer Tonelli®) with a 2-m capacity and precision of 0.1 cm) and weight (measured using a digital scale Camry®, model EB9013, Brazil) with a capacity for 150 kg and a precision of 100 g). Children in the 85th to 97th percentile for age were classified as overweight, and those above the 97th percentile were classified as obese, as recommended by the World Health Organization (WHO)^15^. For the orthodontic assessment, a single experienced orthodontist examined the children^16^, as follows.

### Sagittal analysis

The molar relationship followed Anglés classification^17^, based on the comparison of the anteroposterior diameter of the mandible and maxilla and the relationship between the permanent first molars.

Overjet, the projection of the maxillary incisors beyond the lower incisors on the horizontal plane, measured by using a Community Periodontal Index probe (CPI), placed perpendicular to the occlusal plane, on the buccal surface of the mandibular incisors^18^. A measurement of 0 mm classifies as an edge to edge bite; 1 to 3 mm normal; 4 to 6 mm, moderately increased, and higher than 6 mm, severely increased. In this report, values equal or less than 3 mm, are not classified as overjet; for values higher than are 3mm, classified as overjet. When the maxillary incisors occlude behind the mandibular incisors, negative overjet, then recorded as anterior crossbite ^19,20^.

### Vertical analysis

Overbite is the overlap of the upper teeth along the lower teeth on the vertical plane^16^. The distance (in mm) between the incisal edges, measured using a CPI probe^18^. A measurement of 0 mm (incisal edges of maxillary incisors in contact with incisal edges of mandibular incisors) classifies as an edge to edge ^19,20^; 1 to 3 mm classifies as normal; 4 to 6 mm, moderately increased, and higher than 6mm, severely increased. In the present study, values smaller or equal to 3 mm did not indicate overbite, when higher than 3 mm, record overbite. A lack of contact between the incisal edges of the maxillary and mandibular teeth was considered anterior open bite ^16,19,20^.

### Transversal analysis

Posterior crossbite, defined as a labiolingual relationship between the maxillary and mandibular teeth, when out of the normal standards, Inverted.^16^ this is classified as present, when involving one or more molars in the posterior region. Posterior crossbite, classified as unilateral (involving only one side) or bilateral (involving both sides)^21^.

The orthodontic chart used in this assessment, also meant to record Anglés classification (class I, II or III), overjet, overbite, anterior open bite, anterior crossbite, and posterior crossbite.

The sociodemographics recorded: sex; marital and employment status of the caregiver, and the Sleep Disturbance Scale for Children (SDSC).

SDB was evaluated using the SDSC (for three to 18 years of age), validated for Brazil^22^, and filled by parents or caregiver, about the child’s sleep habits in the previous six months. The scale has 26 items divided among six subscales: Disorders of Initiating and Maintaining Sleep, Sleep Breathing Disorders, Disorders of Arousal, Sleep-Awake Transition Disorders, Disorders of Excessive Somnolence and Sleep Hyperhydrosis. The SDSC items to detect symptoms of SDB were: (1) “the child has difficulty in breathing at night”; (2) “the child gasps for breath or is unable to breathe during sleep”; and (3) “the child snores”. Answers, recorded on a Likert scale of 1–5: 1 point for “never”, 2 points for “occasionally” (once or twice per month), 3 points for “sometimes” (once or twice per week), 4 points for “often” (3–5 times per week), and 5 points for “always” (daily). Hence, the sum score for the three questions can be at least 3 and at most 15. According to the total score obtained for these three SDSC items, participants were classified into 3 groups: group 1, at high risk of SDB (12–15 points); group 2, at moderate risk of SDB (7–11 points); and group 3, at low risk of SDB (3–6 points). This variable, dichotomized in children without SDB (group 1 and 2) and children with SDB (group 3)^16^. This study also evaluated the BMI of children, since according to Kaditis el al^23^ this variable is an important factor for SDB.

Data analysis was performed with the Statistical Package for the Social Sciences (SPSS, version 17.0) by the Pearson’s chi-square test and Lemeshow test, with a 5% margin of error and 95% confidence interval.

## RESULTS

A total of 390 out of the 435 students (89.7%) had physical, and orthodontic examination and parents returned the questionnaires. The losses were 45 students (10.3%) that did not complete the questionnaires or did not attend school on the examination days. The majority of the children was seven years of age (n=212; 54.4%), and males accounted for 53.8% (n=210) of the sample. Among the caregivers, the majority was female (n= 363; 93.1%), married or in a stable relationship (n= 238; 61.0%), who had completed high school or university education (n= 214; 54.9%), n = 242; 62.1% were employed. Table 1 shows to the frequency of malocclusion, body mass index and SDB.

**Table 1.**
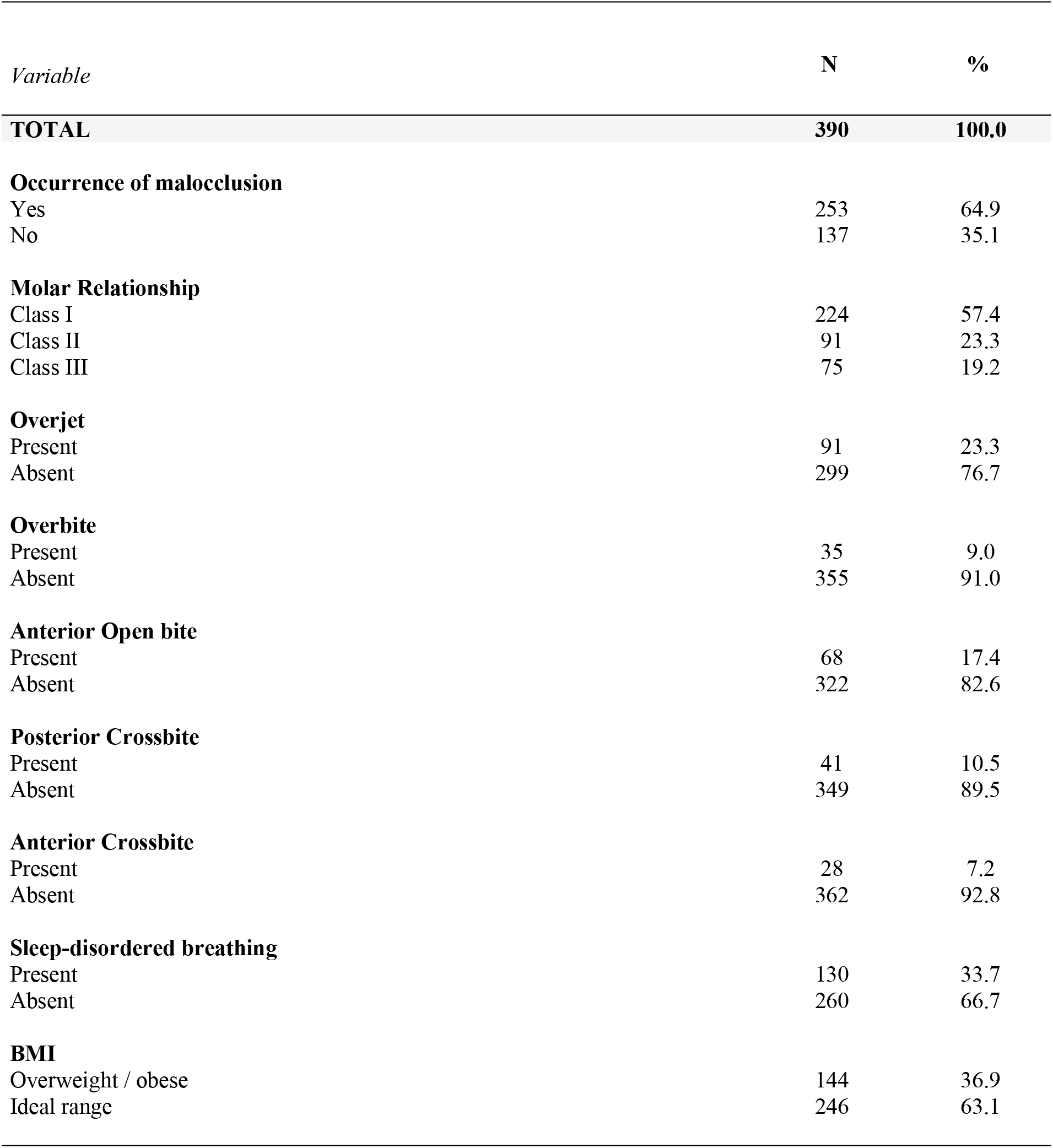
Occurrence of malocclusion, SDB and BMI status in participating children

| *Variable* | **N** | **%** |
| --- | --- | --- |
| **TOTAL** | **390** | **100.0** |
| **Occurrence of malocclusion** |  |  |
| Yes | 253 | 64.9 |
| No | 137 | 35.1 |
| **Molar Relationship** |  |  |
| Class I | 224 | 57.4 |
| Class II | 91 | 23.3 |
| Class III | 75 | 19.2 |
| **Overjet** |  |  |
| Present | 91 | 23.3 |
| Absent | 299 | 76.7 |
| **Overbite** |  |  |
| Present | 35 | 9.0 |
| Absent | 355 | 91.0 |
| **Anterior Open bite** |  |  |
| Present | 68 | 17.4 |
| Absent | 322 | 82.6 |
| **Posterior Crossbite** |  |  |
| Present | 41 | 10.5 |
| Absent | 349 | 89.5 |
| **Anterior Crossbite** |  |  |
| Present | 28 | 7.2 |
| Absent | 362 | 92.8 |
| **Sleep-disordered breathing** |  |  |
| Present | 130 | 33.7 |
| Absent | 260 | 66.7 |
| **BMI** |  |  |
| Overweight / obese | 144 | 36.9 |
| Ideal range | 246 | 63.1 |

Among the orthodontic variables, overjet, anterior, and posterior crossbite were significantly associated with the SDB (TABLE 2). A logistic regression model with the variables that showed a significant association up to 20% (p <0.20) in the bivariate study (overjet, anterior open bite. Posterior crossbite) was used to verify which variables influenced the presence of SDB. In the multivariate logistic regression analysis, posterior crossbite and open bite were considered risk factors for SDB (TABLE 3).

**Table 2.** Occurrence of SDB according to malocclusions and BMI status

| Sleep Breathing Disorders |  |  |  |  |  |  |  |  |
| --- | --- | --- | --- | --- | --- | --- | --- | --- |
| Variable | Present |  | Absent |  | Total |  | p-value | OR (95% CI) |
|  | N | % | n | % | N | % |  |  |
| **Total Group** | **130** | **33.3** | **260** | **66.7** | **390** | **100.0** |  |  |
| **Molar Relationship** |  |  |  |  |  |  | p<sup>(1)</sup> = 0.559 |  |
| Class I | 70 | 31.3 | 154 | 68.8 | 224 | 100.0 |  | 1.00 |
| Class II | 34 | 37.4 | 57 | 62.6 | 91 | 100.0 |  | 1.31 (0.79 a 2.19) |
| Class III | 26 | 34.7 | 49 | 65.3 | 75 | 100.0 |  | 1.17 (0.67 a 2.03) |
| **Overjet** |  |  |  |  |  |  | p<sup>(1)</sup> = 0.007* |  |
| Present | 41 | 45.1 | 50 | 54.9 | 91 | 100.0 |  | 1.93 (1.19 a 3.13) |
| Absent | 89 | 29.8 | 210 | 70.2 | 299 | 100.0 |  | 1.00 |
| **Overbite** |  |  |  |  |  |  | p<sup>(1)</sup> = 0.616 |  |
| Present | 13 | 37.1 | 22 | 62.9 | 35 | 100.0 |  | 1.20 (0.58 a 2.47) |
| Absent | 117 | 33.0 | 238 | 67.0 | 355 | 100.0 |  | 1.00 |
| **Anterior Open bite** |  |  |  |  |  |  | p<sup>(1)</sup> = 0.008* |  |
| Present | 32 | 47.1 | 36 | 52.9 | 68 | 100.0 |  | 2.03 (1.19 a 3.46) |
| Absent | 98 | 30.4 | 224 | 69.6 | 322 | 100.0 |  | 1.00 |
| **Posterior Crossbite** |  |  |  |  |  |  | p<sup>(1)</sup> = 0.001* |  |
| Present | 23 | 56.1 | 18 | 43.9 | 41 | 100.0 |  | 2.89 (1.59 a 5.58) |
| Absent | 107 | 30.7 | 242 | 69.3 | 349 | 100.0 |  | 1.00 |
| **Anterior Crossbite** |  |  |  |  |  |  | p<sup>(1)</sup> = 0.488 |  |
| Present | 11 | 39.3 | 17 | 60.7 | 28 | 100.0 |  | 1.32 (0.60 a 2.91) |
| Absent | 119 | 32.9 | 243 | 67.1 | 362 | 100.0 |  | 1.00 |
| **BMI** |  |  |  |  |  |  | p<sup>(1)</sup> = 0.975 |  |
| Obese | 31 | 34.1 | 60 | 65.9 | 91 | 100.0 |  | 1.05 (0.63 a 1.75) |
| Overweight | 18 | 34.0 | 35 | 66.0 | 53 | 100.0 |  | 1.05 (0.56 a 1.96) |
| Ideal range | 81 | 32.9 | 165 | 67.1 | 246 | 100.0 |  | 1.00 |

(\*) Significant association at 5% level // (1) Pearson's chi-square test

**Table 3.** Multivariate logistic regression analysis of the association between SDB and overjet, anterior open bite and posterior crossbite

| Variable | Bivariate |  | Adjusted |  |
| --- | --- | --- | --- | --- |
|  | OR and 95% CI | p - value | OR and 95% CI | p - value |
| **Overjet** |  | 0.007* |  | 0.402 |
| Present | 1.93 (1.19 a 3.13) |  | 1.29 (0.71 a 2.36) |  |
| Absent | 1.00 |  | 1.00 |  |
| **Anterior Open bite** |  | 0.008* |  | 0.002* |
| Present | 2.03 (1.19 a 3.46) |  | 2.34 (1.36 a 4.03) |  |
| Absent | 1.00 |  | 1.00 |  |
| **Posterior Crossbite** |  | 0.001* |  | 0,014* |
| Present | 2.89 (1.59 a 5.58) |  | 2.79 (1.24 a 6.31) |  |
| Absent | 1.00 |  | 1.00 |  |

(\*) Significant association at 5% level // Lemeshow Test

## DISCUSSION

Among the 390 children, in this study, 65% presented malocclusion; the result is in agreement with previous studies, which report prevalence rates ranging from 60 to 70%. Angle class I was the most prevalent, followed by classes II and III, which is also in agreement with the previous studies ^24,25^., conducted in different countries, which report a high prevalence of malocclusion ^26,27^, that impact the functional, emotional and social aspects of childreńs life^16,28^.

It merits consideration that the present study provides data on the prevalence of malocclusion among children with SDB, which is not a piece of trivial information. According to other studies, untreated SDB in childhood could have a negative impact in adulthood, with an increase in health-related quality of life ^3,6,14,29^. As described in latter reports, Orthodontic intervention in the mixed dentition may prevent or arrest incipient malocclusions, contributing to the harmonious growth of the basal bones and to the normal development of the occlusion, reducing the chances of deleterious disorders in the permanent dentition. ^3,6,14^

SDB may impact public health throughout the world, once it affects millions of individuals, mainly children^30^. SDB often more associated with occlusal abnormalities than other sleep disorders, which may be explained by the abnormal development of malocclusions caused by the breathing pattern, facial muscle balance, and skeletal muscles^30^. This condition can develop into mild to more severe obstructive sleep apnea in a relatively short period in cases of a lack of effective therapy. Adults presenting with SDB seem to have poor dental occlusions and anomalous craniofacial morphology, which may raise a question of whether orthodontic treatment in childhood could prevent SDB in adulthood ^31,32^. Early diagnosis of SDB in childhood and provision of appropriate treatment may not only treat or prevent medium-term complications such as learning difficulties but may potentially prevent long term cardiovascular complications^33^.

Among the occlusal features shown in the present study, overjet, posterior crossbite, and open bite significantly associate with SDB, which is in agreement with data in a previous study ^34^. The logistic regression analysis in the present investigation revealed that both posterior crossbite and open bite were risk factors for sleep-disorders breathing. The present findings indicate the importance of the early diagnosis of posterior crossbite, which seems to be the type of malocclusion most associated with sleep-disordered breathing ^24,31,34^ followed by anterior open bite ^35,36^. Those may lead to an abnormal breathing pattern, which seems to alter the oral and facial muscular balance and are likely to affect skeletal and occlusal development in children^31^.

In this group of children, 36.9% were overweight/obese. The latter is in agreement with previous studies that report a high prevalence of overweight and obesity among children and adolescents aged two to 19 years^31^. Concerning SDB, in this study, there was no significant association with overweight/obese. Likewise, a previous study also found no association between overweight and SDB and, similar to our findings, individuals with craniofacial abnormalities were more susceptible to developing sleep-disorders breathing than individuals with excess weight^37^.

Evidence suggests that obesity plays an important role in the pathophysiology of SDB in adults^38^ and, to a lesser extent, in children^39^. Thus, more knowledge about the interactions between obesity and SDB is relevant when considering the increases in the rates of obesity in childhood ^23,40^.

SDB represents a continuum of manifestations from simple snoring to Obstructive sleep apnea (OSA), and this condition tends to progress from mild SDB to severe OSA over a varying period, which may be surprisingly short in the case of weight gain and the lack of effective treatment ^31,41^

In children, adenotonsillar hypertrophy, as well as deviations concerning craniofacial morphology and dental occlusion, are risk factors for SDB, which seems to be complex, multifactor pathogenesis in children ^31,37^. Thus, timely orthodontic intervention could prevent SDB in adulthood ^11,16^.

The main limitation of the present report is the study design, which does not allow the determination of cause-and-effect relationships between malocclusion and sleep-disorders breathing. Thus, longitudinal design studies are needed for more precise diagnosis purposes.

## CONCLUSIONS

In the study population, the prevalence of SDB was high and associated with malocclusion. Posterior crossbite and anterior open bite were found to be risk factors for SDB.

## Acknowledgements

The authors of this research are grateful to the children and their families who have agreed to participate and CAPES, Brazilian Ministry of Education, for funding.

## Contributions

MCA Lyra: conceptualization, data curation, writing (original draft preparation) and writing (review & editing) of the manuscript; D Aguiar, MC Paiva, M Arnaud and A Vasconcelos: data curation and visualization; A Rosenblatt, NPT Innes and MV Heimer: conceptualization, supervision and writing (review & editing). All authors read and approved the final manuscript and agree to be accountable for it.

